# A novel order-level lineage of ammonia-oxidizing *Thaumarchaeota* is widespread in marine and terrestrial environments

**DOI:** 10.1101/2023.02.17.529030

**Authors:** Yue Zheng, Baozhan Wang, Ping Gao, Yiyan Yang, Xiaoquan Su, Daliang Ning, Qing Tao, Feng Zhao, Dazhi Wang, Yao Zhang, Meng Li, Mari-K.H. Winkler, Anitra E. Ingalls, Jizhong Zhou, Chuanlun Zhang, David A. Stahl, Jiandong Jiang, Willm Martens-Habbena, Wei Qin

**Affiliations:** State Key Laboratory of Marine Environmental Science and College of the Environment and Ecology, Xiamen University, Xiamen, China; Department of Microbiology, College of Life Sciences, Nanjing Agricultural University, Key Laboratory of Agricultural and Environmental Microbiology, Ministry of Agriculture and Rural Affairs, Nanjing, China; National Library of Medicine, National Institutes of Health, Bethesda, MD; College of Computer Science and Technology, Qingdao University, Qingdao, China; Department of Microbiology and Plant Biology and Institute for Environmental Genomics, University of Oklahoma, Norman, OK, USA; CAS Key Laboratory of Urban Pollutant Conversion, Institute of Urban Environment, Chinese Academy of Sciences, Xiamen, China; State Key Laboratory of Marine Environmental Science and College of Ocean and Earth Sciences, Xiamen University, Xiamen, China; Archaeal Biology Center, Institute for Advanced Study, Shenzhen University, Shenzhen, China; Department of Civil and Environmental Engineering, University of Washington, Seattle, WA, USA; School of Oceanography, University of Washington, Seattle, WA, USA; School of Civil Engineering and Environmental Sciences, University of Oklahoma, Norman, OK, USA; Earth and Environmental Sciences, Lawrence Berkeley National Laboratory, Berkeley, CA, USA; Shenzhen Key Laboratory of Marine Archaea Geo-Omics, Department of Ocean Science and Engineering, Southern University of Science and Technology, Shenzhen, China; Southern Marine Science and Engineering Guangdong Laboratory (Guangzhou), Guangzhou, 511458, China; Department of Microbiology and Cell Science & Fort Lauderdale Research and Education Center, University of Florida, Davie, FL, USA

## Abstract

Ammonia-oxidizing archaea (AOA) are among the most ubiquitous and abundant groups of Archaea on Earth, widely distributed in marine, terrestrial, and geothermal ecosystems. However, the genomic diversity, biogeography, and evolutionary process of AOA populations in subsurface environments are vastly understudied compared to those of marine and soil AOA. We here report a novel AOA order *Candidatus* Nitrosomirales that forms a deeply branching basal sister lineage to the thermophilic *Ca.* Nitrosocaldales. Metagenomic and 16S rRNA gene read mapping demonstrates the dominant presence of *Nitrosomirales* AOA in various groundwater environments and their widespread distribution across a range of geothermal, terrestrial, and marine habitats. Notably, terrestrial *Nitrosomirales* AOA show the genetic capacity of using formate as an alternative source of reductant and appear to have acquired key metabolic genes and operons from other mesophilic populations via horizontal gene transfer, including the genes encoding urease, nitrite reductase, and V-type ATPase. Potential metabolic versatility and acquired functions may facilitate their radiation into a variety of subsurface, marine, and soil environments. Molecular thermometer-based evolutionary analysis suggests that *Nitrosomirales* originated from thermophilic environments and transitioned into temperate habitats in parallel with *Nitrososphaerales* and *Nitrosopumilales*. We also provide evidence that terrestrial-marine habitat transitions occurred within each one of the four AOA orders, which reveals a more complex evolutionary trajectory of major AOA lineages than previously proposed. Together, these findings establish a robust taxonomic and evolutionary framework of AOA and provide new insights into the ecology and evolution of this globally abundant functional guild.

## Introduction

The ammonia-oxidizing archaea (AOA) represent one of the most abundant and ubiquitous archaeal groups in the global biosphere [1, 2, 3]. They account for nearly 30% of microbial plankton in the oceans and up to 5% of microbial populations in soils [4, 5]. Apart from their generally high abundance in a vast range of marine and terrestrial environments, they also play an important role in the global nitrogen cycle. They are almost exclusively responsible for ammonia oxidation in oligotrophic marine environments and contribute as much as 80% of the emission of ozone-depleting potent greenhouse gas nitrous oxide (N_2_O) from the oceans [6, 7]. This globally abundant and ecologically significant group of archaea was assigned to a major archaeal phylum *Thaumarchaeota* (also named as *Nitrososphaerota*) [8, 9, 10].

Although all AOA are united by a common physiology of chemoautotrophic growth on ammonia oxidation and carbon fixation, the extensive genetic repertoire of the pan-genome suggests many additional adaptive features are associated with the remarkable ecological success of this functional guild as reflected by their high global abundance and wide niche breadth [11, 12, 13, 14, 15, 16, 17]. Previous AOA phylogenetic, ecological, and evolutionary analyses were based on a standardized taxonomic framework that divided ammonia-oxidizing *Thaumarchaeota* into four major lineages, the *Nitrosopumilales* (Group 1.1 a) [18], *Candidatus* (*Ca*.) Nitrosotaleales (Group 1.1 a-associated, now reclassified as a family within the *Nitrosopumilales*) [19], *Nitrososphaerales* (Group 1.1 b) [20], and *Ca.* Nitrosocaldales (Thermophilic AOA, ThAOA) [21], that appear somewhat specialized, respectively, to aquatic (marine or freshwater), acidic soil, neutral or alkaline soil, and geothermal environments [12, 13, 16, 17, 22, 23]. Intriguingly, a recent metagenomic study of cold deep seawater (∼ 2.3 °C) recovered from the Mariana Trench yielded a metagenome-assembled genome (MAG) that appeared to be phylogenetically closely associated with the deeply branching thermophilic AOA [24]. This marine AOA MAG, along with two other closely related MAGs recovered from deep-sea waters (UBA213 and SAT137), had been assigned to *Ca*. Nitrosocaldales based on the previously established phylogenetic backbone of AOA [11, 12]. However, further phylogenomic analyses with additional genomes that represent a broader range of genotypes of this early branching lineage are needed to resolve and substantiate the uncertain taxonomic affiliation and evolutionary trajectory of basal AOA clades. In addition, how widespread these early branching mesophilic AOA are in marine and terrestrial environments, and their diversity, metabolic adaptation, and ecological significance, remain unknown.

Here we conduct phylogenomic and comparative genomic analyses of 118 AOA and non-ammonia-oxidizing *Thaumarchaeota* genomes, including 23 MAGs and single amplified genomes (SAGs) of this under-studied group that were obtained from a variety of subsurface, geothermal, soil, and marine environments. We show that these 23 MAGs and SAGs form a highly supported monophyletic order-level lineage within the *Nitrososphaeria*, which we here designate a new AOA order *Ca*. Nitrosomirales. The global distribution of *Ca*. Nitrosomirales AOA was investigated by extensively searching for their 16S rRNA and *amoA* gene sequences in marine and groundwater metagenome datasets as well as in the Microbiome Search Engine 2 database that contains over 300,000 samples sequenced from a vast range of terrestrial and aquatic environments [25]. Our findings provide new understanding of the metabolic potential, evolution, and biogeography of this basal branching lineage that appears to represent a predominant AOA genotype in previously under-sampled habitats, such as many terrestrial subsurface environments and deep-sea sponges.

## Materials and Methods

### The distribution of *Ca.* Nitrosomirales AOA at global scale

The 16S rRNA gene sequences of all AOA were extracted by the ssu_finder of CheckM (version 1.0.12) [26]. The extracted AOA sequences were subsequently mapped into the database of Microbiome Search Engine 2 (MSE2, http://mse.ac.cn) as previously described [25]. The MSE2 database contains more than 300,000 16S rRNA gene amplicon and metagenomic sequencing samples collected from marine and terrestrial habitats as well as human, animal and plant associated microbiomes. AOA-containing microbiomes in MSE2 were identified via VSEARCH and using a 97% sequence similarity cutoff, by comparing the amplicon and metagenomic sequences to 16S rRNA sequences extracted from 106 AOA genomes (Table S1) [27]. The relative abundance of AOA amplicon sequences in each sample was normalized using the Meta-Storms algorithm [28] to reduce the 16S rRNA gene amplification bias.

In addition to searching against the 16S rRNA gene amplicon database, the metagenomic data of seawater and groundwater samples were searched against the *amoA* genes (encoding the alpha subunit of ammonia monooxygenase). To explore the distribution of *Nitrosomirales* AOA in the global ocean, we searched *amoA* gene reads from global ocean metagenomic databases, including *Tara*-Oceans (2009-2013) [29] and *Malaspina*-2010 metagenomic data [30], Hawaii Ocean time series station metagenomic data [31], and the Mariana Trench metagenomic data [32]. To confirm the prevalence of *Nitrosomirales* AOA in the contaminated groundwaters, we also searched *amoA* gene reads from the contaminated groundwater metagenomic samples collected from the Oak Ridge Integrated Field Research Challenge (OR-IFRC) experimental sites (FW300, FW301, FW602, and GW715) [33]. Briefly, we first trimmed the raw sequencing reads of metagenomic samples by Trimmomatic (version 0.36) [34]. The trimmed reads were then mapped via Diamond (version 0.9.24.125) [35] with a threshold of 80% similarity and 100 bp coverage, as well as an E-value cut-off of 1 × e^-10^, to an in-house AOA species genome database containing *amoA* genes found in available AOA species, MAGs, and SAGs. The recruited reads were further classified into different AOA lineages. In addition, we searched for 16S rRNA and *amoA* gene sequences of *Nitrosomirales* in the NCBI database with BLASTN using a sequence similarity cutoff of 95% and 90%, respectively, and an E-value cut-off of 1 × e^-10^. The phylogenetic affiliation of the identified sequences was further confirmed in the 16S rRNA and *amoA* gene trees. Only the sequences affiliated by a bootstrap value of more than 85% (out of 1000 replications) within the *Nitrosomirales* clade were identified as the members of *Nitrosomirales*.

### Comparative genomic analysis and pathway construction

Average nucleotide identity (ANI) was calculated by pyani (https://github.com/widdowquinn/pyani) using the model of BLASTN. The percentage identity and alignment coverage between each of two genomes were displayed in heatmap format. The corresponding genome size and GC content were marked for each AOA genome. AOA proteins were clustered into homolog cluster groups (HCGs) by OrthoMCL (version 2.0.9) [36]. The thresholds of HCGs were set to achieve a pairwise coverage of 50% and identity of 50% based on all-against-all BLASTP. Orthologs and paralogs were identified as the reciprocal best similarity pairs that were found between and within species, respectively [37]. Core genome represents the HCG genes shared by all AOA species genomes, MAGs, and SAGs, and pan-genome represents the collective set of genes present in at least one AOA genome. For *m* genomes selected out of n strains, a total of *n*!/ (*m*-1)!·(*n-m*)! combinations were calculated to determine the sizes of the core and pan-genomes. To compare the pan-genome openness of different AOA orders, the average number of new unique genes per Mbp genome was calculated with the sequential addition of each AOA genome. Up to 5,000 random combinations were sampled for core genome and pan-genome analyses. Finally, the pan-genomes of *Nitrosomirales* and other AOA orders were visualized using the Anvi’o software (version 2.2.2) [38].

The functions of HCGs were subsequently annotated according to the reference AOA genomes (*Ca.* Nitrosocaldus cavascurensis SCU2 [39] and *Nitrosopumilus maritimus* SCM1 [40]). The putative metabolic pathways were classified into 16 groups, including ammonia oxidation/assimilation/nitrate reduction, urea utilization, carbon fixation/metabolism, sulfur assimilation/metabolism, phosphorus utilization, stress response, CPISPR system, thermo-adaptation, amino acids/vitamins/cofactors, information processes, S-layer synthesis, lipid biosynthesis, glycosyl transferase, hydrogen oxidation, ectoine synthesis, and transporters. Putative transporters were further identified by screening against the Transporter Classification Database [41].

### Optimal growth temperature estimates for *Nitrosomirales* AOA

The stem regions of *Nitrosomirales* AOA 16S rRNA genes were predicted based on compensatory base-pair changes using the RNAfold package [42], and the GC contents of the predicted stem were calculated for the *Nitrosomirales* genome-retrieved 16S rRNA gene sequences. The evolutionary model of AOA 16S rRNA sequences along the reference tree was estimated by a nonhomogeneous model using bppml, and a total of 100 replicates of ancestral sequences were constructed using the program BppAncestor (BppSuite version 2.4.1) as previously described [13, 43]. The optimal growth temperatures (OGTs) of the ancestors of *Nitrosomirales* AOA were estimated using the previously calibrated linear regression between the GC contents of AOA 16S rRNA gene stem regions and their OGTs [13]. The OGTs with confidence intervals of corresponding nodes (ancestor nodes of all AOA and the four orders) were inferred against the linear regression.

### Phylogenomic tree construction

Phylogenomic tree construction was based on concatenated alignments of 70 conserved marker genes (Table S2) from 106 AOA genomes (Table S1) and outgroup archaeal species. The genomes (*Nitrosomirales sp*. UBA213 and *Nitrosomirales sp*. WS1) that had not been annotated in NCBI and JGI were annotated using GeneMarks in this study [44] (Supplementary Datasets S1 and S2). The outgroup species for phylogenomic analysis included non-ammonia-oxidizing *Thaumarchaeota*, *Aigarchaeota*, *Bathyarchaeota*, *Crenarchaeota*, *Korarchaeota*, *Lokiarchaeota*, *Euryarchaeota*, and *DPANN* archaeal species. The 70 marker genes were identified by BLASTP, and they were individually aligned by MAFFT (version 7.221) [45]. Subsequently, the conserved regions of alignments were extracted by Gblocks (version 0.91b) [46]. Conserved regions were concatenated as a single evolutionary unit. For phylogenomic analysis, the best-fit model of amino acid substitution was estimated using ProtTest (version 3.4) [47]. Maximum likelihood phylogenomic trees were built with concatenated sequences via RAxML (version 8.0.26) using the LG+I+G+X model on the basis of 100 bootstrap replications [48].

### Phylogenetic tree of key genes

The sequences of 16S rRNA, *amoA*, *atpA* and *atpC* (encoding the alpha and epsilon subunits of ATPase, respectively), *nirK* (encoding putative nitrite reductase), and *ureC* genes (encoding the alpha subunit of urease) were extracted from the collected 106 AOA genomes (Table S1). Additional *Nitrosomirales* 16S rRNA genes and 33,387 AOA *amoA* genes were collected from NCBI and extracted from a previous *amoA* gene-based phylogenetic analysis [22], respectively, for comparative phylogenetic analysis.

The phylogenetic trees of the 16S rRNA and other key functional genes of AOA were built using IQ-TREE (v. 2.1.2 COVID-edition) with best-fit model selection [49]. Gene sequence alignment was carried out using MAFFT (v7.407) [45] and was then edited with Gblocks (v0.91b) [50] to identify conserved regions. The best-fit model of evolution for each gene set was selected by ModelFinder [51] with “-m MF –T AUTO” flag as follows: GTR+F+R4 for 16S rRNA genes, GTR+F+I+Γ4 for *amoA* genes, LG+R5 for *atpA*, LG+R4 for *atpC* genes, WAG+F+G4 for *nirK* genes, and LG+R3 for *ureC* genes. Subsequently, the maximum likelihood phylogenetic trees were built using IQ-TREE with best model mentioned above and 1,000 times ultrafast bootstrapping [52]. The non-ammonia-oxidizing *Thaumarchaeota* were used as the outgroup for the 16S rRNA gene tree. *Thermoproteales*, *Sulfolobus*, *Desulfurococcales Ignicoccus hospitalis* KIN4/I, *Ca*. Bathyarchaeota archaeon ex4484 231, and *Methanosuratus*/*Methanomethylicus* were used as the outgroups for the phylogenetic trees of the subunits A and C of the A-type ATPases, and *Thermoplasmatales* as well as *Enterococcus hirae* ATCC 9790 were used as the outgroups for the trees of the subunits A and C of the V-type ATPases.

### Availability of data and materials

AOA genomes used in this study are available in the NCBI and JGI databases. The corresponding accession numbers are listed in Table S1. The annotation file of *Nitrosomirales* MAGs UBA213 and WS1 are provided in GFF format (Supplementary Datasets S1 and S2). Completeness, contamination, and coding density of AOA genomes were assessed by CheckM (version 1.0.12) [53]. All other data products associated with this study are available from the corresponding authors upon request.

## Results and Discussion

### *Nitrosomirales* order represents a basal lineage of AOA

We compiled a 118-genome dataset that comprises 106 AOA cultured genomes, MAGs, and SAGs as well as 12 representative genomes of non-ammonia-oxidizing *Thaumarchaeota* to serve as outgroups (Table S1). To infer the most probable evolutionary relationship among AOA taxa and resolve the uncertain taxonomic affiliation of mesophilic members in the deepest branching AOA lineages, we constructed a maximum likelihood phylogenomic tree from the concatenation of 70 conserved marker genes (Table S2) present in AOA genomes and outgroup thaumarchaeotal MAGs. We obtained 20 additional MAGs and SAGs from various terrestrial subsurface, deep soil, and marine habitats that were clustered with three previously reported marine AOA MAGs MTA5, UBA213, and SAT137 [11, 12, 24], and they together formed a well-supported monophyletic group branching as a sister clade to the thermophilic *Ca*. Nitrosocaldales AOA (Fig. 1). The monophyletic phylogeny of this deeply branching lineage did not emerge in previous analyses using single-gene markers or only with the three reported marine AOA MAGs [11, 12, 16, 24]. Among these newly obtained MAGs and SAGs, nine were retrieved from **w**arm/thermal **s**pring (WS1–WS3) and **c**arbonate **s**pring (CS1–CS6) waters and sediments, six originated from groundwaters (GW1–GW6), two from **b**asaltic **l**ava **c**aves (BLC1 and BLC2), two from **d**eep **s**andy **s**oils in switchgrass fields (DSS1 and DSS2), and one from **d**eep **o**cean waters (OD1) (Table S3). Eleven of these genomes showed a generally high level of completeness (> 78%) and low contamination (< 5%) (Table S3). MAGs DSS1 and WS1 are complete or nearly complete genomes and estimated to be 100% and 99.93% complete with minimal contamination (0.97%), respectively. These MAGs and SAGs span a large size range from 0.99–3.68 Mbp, and similar to other AOA genomes, the GC content of marine *Nitrosomirales* genomes (34.8–36.1%) was lower than that of terrestrial *Nitrosomirales* genomes (40.3–47.7%) (Table S3). The average nucleotide identity (ANI) values among these 23 genomes were 67.6%–99.6% (Table S4), reflecting generally high genomic diversity within this group. The characterized *Nitrosomirales* genomes shared low genomic homology (<68.2% ANI) and alignment fraction (4.8%–29.4%.) with other marine and terrestrial AOA genomes (Table S4).

**Figure 1.**
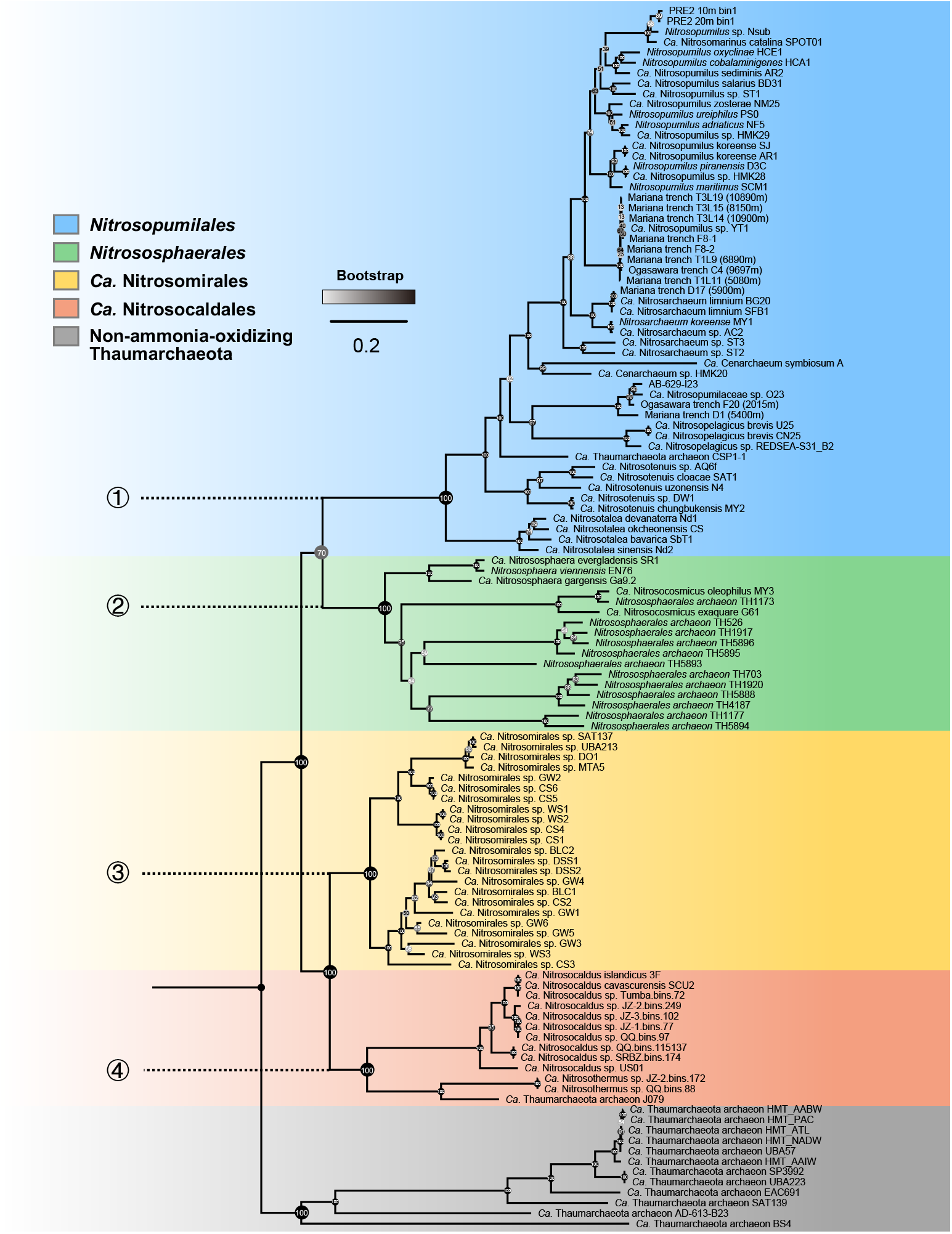
Phylogeny of *Ca*. Nitrosomirales and other AOA orders. Phylogenomic inference of AOA species affiliated to the orders *Nitrosopumilales* (blue, basal lineage #1), *Nitrososphaerales* (green, basal lineage #2), *Ca*. Nitrosomirales (yellow, basal lineage #3), and *Ca*. Nitrosocaldales (orange, basal lineage #4) based on concatenated sequences of 70 conserved single-copy core genes. The non-ammonia-oxidizing thaumarchaeotal genomes were used as outgroups (grey). Please note that, based on the recent comparative phylogenetic analysis of AOA genomes [12], the previously defined acidophilic AOA group *Ca.* Nitrosotaleales has been merged with *Nitrosopumilales*. Confidence values were provided based on 100 bootstrap replications.

Additional whole-genome based taxonomic analysis using the Genome Taxonomy Database Toolkit (GTDB-Tk) also showed these 23 MAGs and SAGs to be members of a distinct basal lineage along with the other three formally described major AOA lineages (Table S3) [18, 20, 21]. We here propose the name *Ca.* Nitrosomirales to represent this deeply branching AOA order within the class *Nitrososphaeria* (Fig. 1), branching between the orders *Nitrososphaerales* and thermophilic *Ca*. Nitrosocaldales with the complete MAG DSS1 from deep sandy soil representing the candidate family *Nitrosomiraceae* and genus *Nitrosomirus*. The name *Nitrosomirus* refers to the organisms as ammonia oxidizers with the capacity of oxidizing ammonia to nitrite (from the Latin “*nitrosus”*, full of natron; here intended to mean nitrous) and its surprisingly wide distribution range in various marine, soil, geothermal, and subsurface habitats (from the Latin adjective “*mirus*”, meaning surprising and amazing), spanning a range of temperature, salinity, pressure, and nutrient availability (see the description below).

### *Nitrosomirales* AOA are widely distributed in diverse terrestrial, marine, and geothermal habitats

The recovery of many *Nitrosomirales* MAGs from subsurface, deep soil, deep ocean, and geothermal environments suggest that *Nitrosomirales*-AOA are ubiquitous in diverse terrestrial and marine habitats. To further explore the distribution of this under studied group in the global biosphere, we searched the *Nitrosomirales* MAG-derived 16S rRNA sequences against the Microbiome Search Engine 2 16S rRNA gene database that contains over 300,000 16S rRNA gene amplicon and metagenomic sequencing samples obtained from a broader range of natural habitats and engineered systems, as well as human, animal and plant associated microbiomes (see Materials and Methods for details) [25]. *Nitrosomirales*-derived 16S rRNA gene sequences were found in various terrestrial and marine habitats, including groundwaters, farmland soils, forest soils, wetlands, and lake sediments, marine sponges, and marine aquaculture waters (Figs. 2 and Table S5). Additional extensive searches of 16S rRNA (similarity cutoff of 95% and E-value cutoff of 1 × e^-10^) in NCBI database further identified *Nitrosomirales* sequences in (moderate)thermophilic habitats, such as hydrothermal vents (>52.0 °C and 72–103 °C), hot springs (55.0 °C), and thermal karst well waters (73.7 °C) (Fig. 2). Both the 16S rRNA and *amoA* gene-based phylogenies support the monophyletic grouping of *Nitrosomirales* metagenomic, amplicon, and clone sequences (Figs. 2 and S1). Together, our results indicate that the habitats of *Nitrosomirales* AOA include a wide variety of terrestrial, marine, subsurface, and geothermal environments (Fig. 3a). Thus, the extraordinary habitat range of *Nitrosomirales* exceeds that of any previously described AOA order-level lineages.

**Figure 2.**
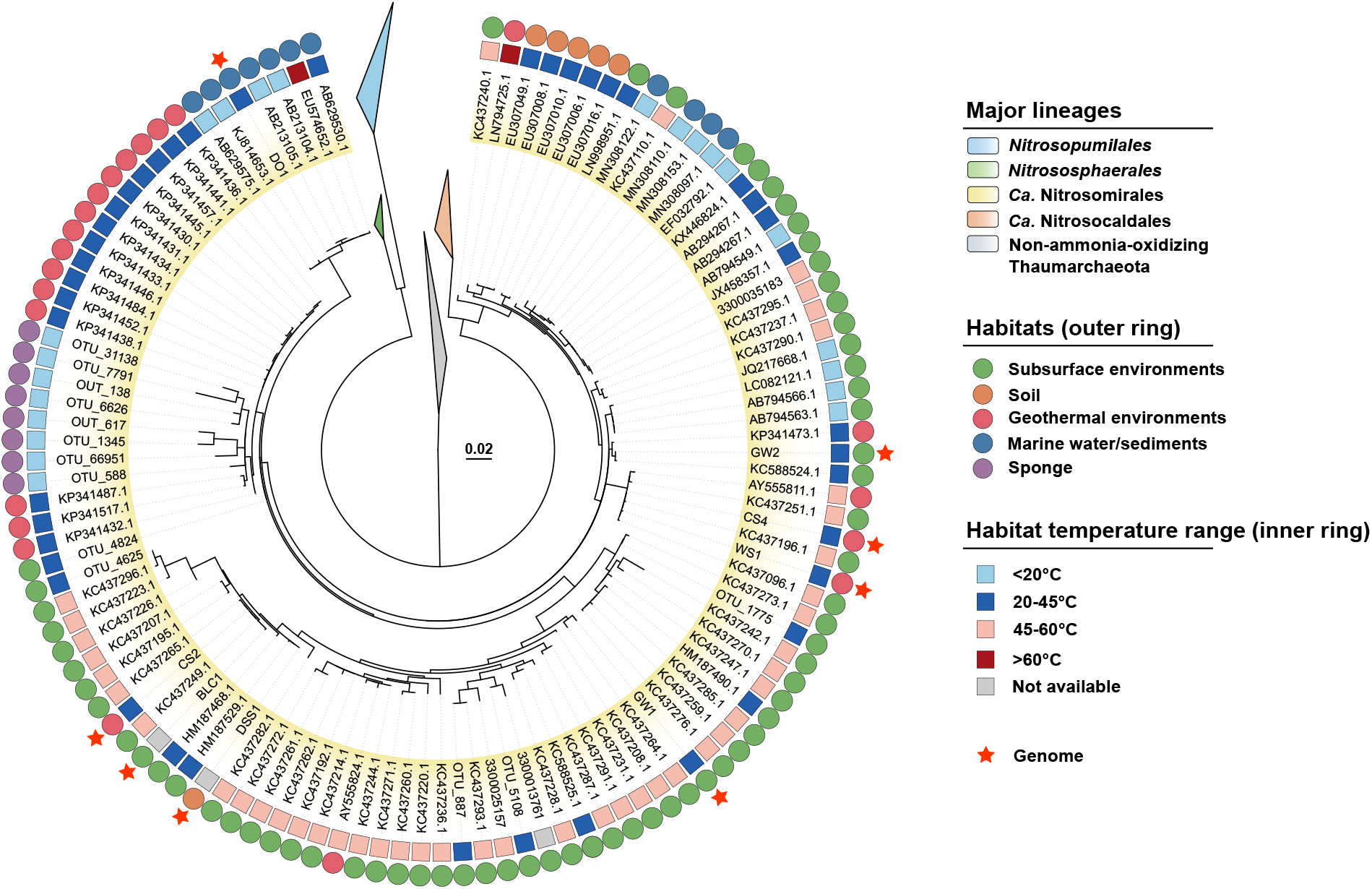
Phylogenic distribution of *Ca*. Nitrosomirales 16S rRNA genes according to habitat types (squares, inner ring) and temperature ranges (circles, outer ring). Confidence values are on the basis of 1000 bootstrap replications. The scale bar represents 2% estimated sequence divergence. The 16S rRNA sequences that retrieved from *Ca*. Nitrosomirales genomes were indicated with stars. The 16S rRNA genes of other thaumarchaeotal lineages were collapsed as triangles.

**Figure 3.**
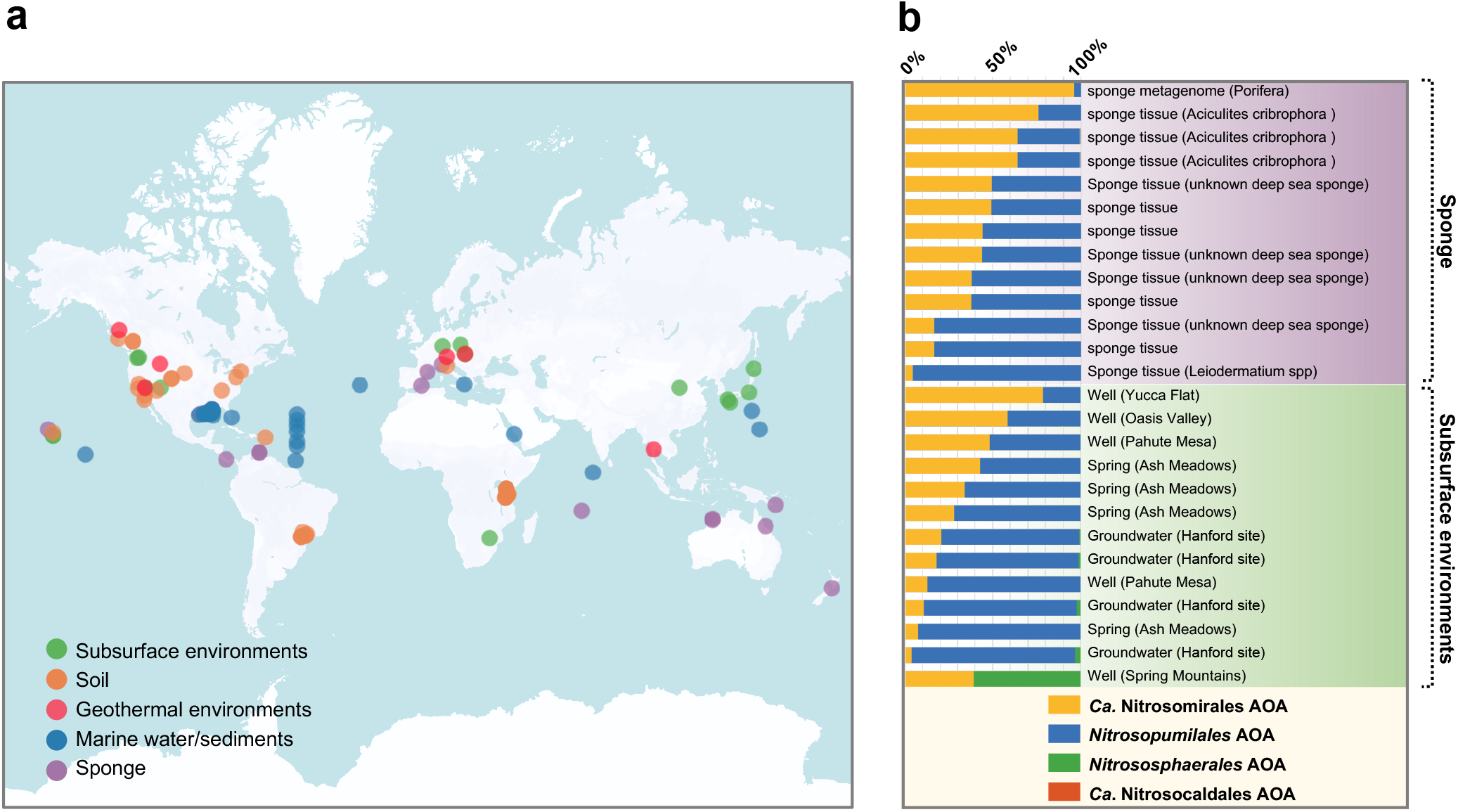
The global distribution of *Nitrosomirales* AOA in various terrestrial and marine environments (**a**). *Nitrosomirales* AOA were enriched in sponge-associated microbiomes and subsurface environments (**b**). Relative abundance of *Nitrosomirales* AOA to total AOA were estimated by comparing the number of amplicon sequencing and metagenomic reads mapping to the 16S rRNA and *amoA* genes of *Nitrosomirales* AOA relative to those of total AOA in the Microbiome Search Engine 2 global 16S rRNA database, groundwater metagenomes and global ocean metagenome databases.

### *Nitrosomirales* AOA represent the dominant AOA population in subsurface habitats and deep-sea sponges

Although the first three *Nitrosomirales* MAGs were recovered from the dark ocean, *Nitrosomirales* AOA were rarely detected in global ocean metagenomes. We assessed the number of metagenomic reads mapping to the *amoA* genes of *Nitrosomirales* AOA and other marine AOA genotypes in global ocean metagenome databases across four oceans and two seas, spanning from epipelagic to hadopelagic zones [29, 30, 31, 32], and found that *Nitrosomirales* AOA only comprised at most 0.5% of the total AOA community in well-studied ocean waters (Table S5). In contrast, the 16S rRNA and *amoA* gene read recruitment showed that *Nitrosomirales* AOA can represent an abundant AOA genotype or even exclusively represent the whole AOA community in some previously under-sampled habitats, such as deep-sea sponges and groundwaters (Fig. 3b and Table S5).

Marine AOA have been frequently reported to associate with sponges [54, 55, 56, 57, 58] and often dominate the whole archaeal communities of sponge microbiomes [59]. Since the sequencing of the first sponge AOA symbiont *Ca.* Cenarchaeum symbiosum hosted by the demosponge *Axinella mexicana* [54], additional AOA MAGs were obtained from various shallow-water and deep-sea sponge-associated microbiomes. Marine sponges are a highly diverse clade of metazoans that contains 125 families, 680 genera, and 11,000 species [60]. The molecular surveys of sponge microbiota indicated that microbial communities were mostly specific to sponge species [61], and the dominant AOA populations were even specific to individual sponges [58]. Previous studies of thaumarchaeotal sponge symbionts only focused on several limited sponge families of the classes *Demospongiae* and *Hexactinellida*. These sponge symbionts were affiliated with five different AOA genera, *Nitrosopumilus* [56], *Ca*. Cenarchaeum [54], *Ca.* Nitrosopelagicus [58], *Ca.* Nitrosospongia [55], and *Ca.* Cenporiarchaeum [56], all of which were within the order *Nitrosopumilales*.

We found that, apart from *Nitrosopumilales*, the members belonging to *Nitrosomirales* can also constitute a significant fraction (up to 96.05%) of the total AOA populations in deep-sea sponges (Fig. 3b). *Nitrosomirales* AOA were specifically hosted by the deep-sea (∼200–550 m) *Aciculites* sponge species within the *Scleritodermidae* family and the *Porifera* phylum (Table S5), and this lineage of sponges was poorly represented in the previous 16S rRNA gene surveys and the global sponge microbiome metagenomic database. Intriguingly, the 16S rRNA gene sequences that are closely related to *Ca*. Nitrosocosmicus AOA within the *Nitrososphaerales* were found in marine sponges *Spirastrella panis* [62], *Astrosclera willeyana* [63], *Theonella swinhoei* [64], *Pseudoceratina purpurea* (NCBI accession No.: KU064739), and *Halichondria oshoro* (HM101091) associated microbiomes (Fig. S2). These findings significantly expand the genetic diversity of the sponge-associated marine AOA beyond the order *Nitrosopumilales*. It is very likely that *Nitrosomirales* AOA play an important role in the nitrogen metabolism and nitrogenous waste removal of deep-sea *Aciculites* sponges, similar to other characterized sponge-associated *Nitrosopumilales* AOA [54, 55, 56].

Another relatively under-sampled AOA habitat is terrestrial subsurface environment. The biogeography of AOA in groundwater and cave ecosystems and associated environmental variables that control the abundance and composition of AOA communities in these systems are poorly documented [65]. Given that several *Nitrosomirales* MAGs were retrieved from the Death Valley Regional Flow System (DVRFS), we leveraged the available 16S rRNA amplicon sequencing data [66] collected from the DVRFS region to assess the relative abundance of *Nitrosomirales* AOA across three major groundwater basins (Table S5). *Nitrosomirales* AOA 16S rRNA genes were detected in nine DVRFS groundwater sites out of the 36 total sampling sites. Based on 16S rRNA gene read recruitment, we found that *Nitrosomirales* AOA can constitute a substantial proportion (30.94–100.00%) of the total AOA populations in six of these groundwater sites (Fig. 3b and Table S5). *Nitrosomirales* AOA were detected in both shallow (0–25 m sampling depth) and deep (474–700 m sampling depth) aquifers with distinct aquatic geochemistry, including Ca-Mg-HCO_3_, Na-HCO_3_, and NaCl dominated groundwaters (Table S5). In addition, they were also found in basaltic lava caves (Table S3), together indicating their wide distribution in terrestrial subsurface environments.

Interestingly, we also identified this lineage in the 16S rRNA gene and metagenome datasets collected from the groundwaters at the Hanford site and the Oak Ridge Integrated Field Research Challenge (OR-IFRC) site, respectively (Fig. 3b and Table S5), legacies of the Manhattan Project contaminated with mixed waste, including radionuclides and nitrate [67]. *Nitrosomirales* AOA were not detected in heavily uranium-contaminated groundwaters but can be enriched in groundwaters with relatively high nitrate concentrations (as high as 816.56 mg/L). It is conceivable that respiratory ammonifiers with the capacity of the dissimilatory nitrate reduction to ammonium (DNRA) could supply ammonia to *Nitrosomirales* AOA and other archaeal and bacterial nitrifiers in these contaminated groundwater sites [68], and thus together these microbial communities may contribute to nitrogen transformation in terrestrial subsurface ecosystems.

### Genomic features and metabolic potential of *Nitrosomirales* AOA

We calculated the core genome and pan-genome of *Nitrosomirales* and other AOA orders to get quantitative insights into the conserved and flexible gene pools of ammonia-oxidizing *Thaumarchaeota* (Fig. S3). Comparative genomic analysis showed that all terrestrial and marine *Nitrosomirales* AOA MAGs and SAGs harbored the conserved pathway genes that are involved in the characterized central metabolism of AOA, including the copper-dependent respiration and electron transfer systems, 3-hydroxypropionate/4-hydroxybutyrate autotrophic carbon fixation cycle, and the biosynthesis of the B vitamin cofactors thiamin (B_1_), riboflavin (B_2_), pantothenate (B_5_), pyridoxine (B_6_), biotin (B_7_), and cobalamin (B_12_) (Fig. 4 and Table S6). The mesophilic *Nitrosomirales* AOA also contained many genes that were thought to be present only in thermophilic *Ca*. Nitrosocaldus/Nitrosothermus AOA and associated with compatible solutes production for thermoprotection [39, 69], such as the genes encoding mannosyl-3-phosphoglycerate synthase and cyclic 2, 3-diphosphoglycerate synthetase (Table S6). This may be due to the retention of ancestral features prior to the adaptive divergence of AOA into mesophilic habitats or may suggest completely different functions of these compatible solutes in *Nitrosomirales* and *Ca*. Nitrosocaldales AOA.

**Figure 4.**
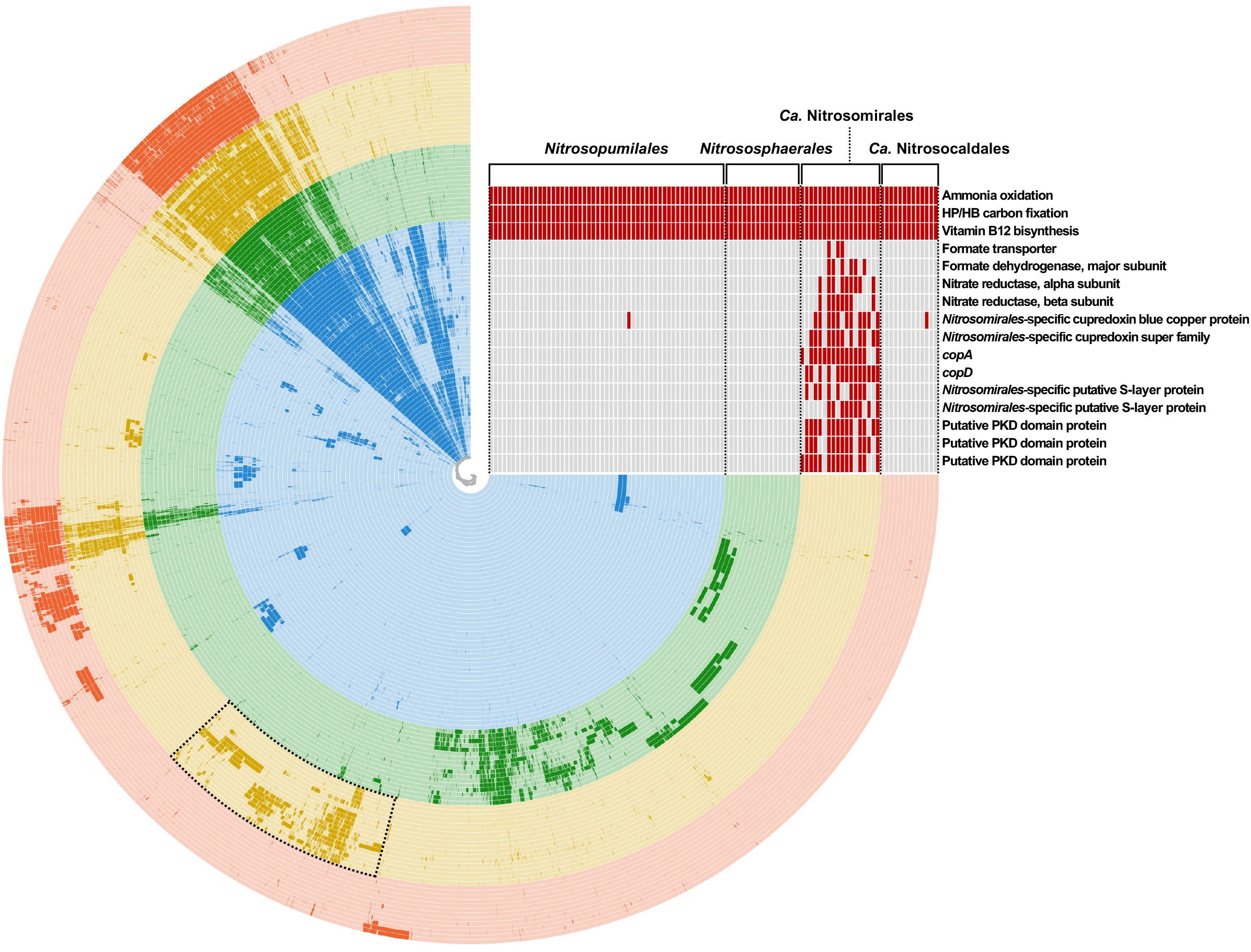
Anvi’o representation of the pangenome of *Nitrosomirales* and other AOA genomes. Each radial layer represents an AOA genome, which was arranged by the order of phylogenomic tree shown in Fig. 1. In each layer, dark and light colors represent the presence and absence of protein clusters, respectively. The heatmap in the top right corner represents the conserved functions present in the core genome of all AOA orders and highlights key unique gene contents retrieved from the protein clusters that are specifically enriched in *Nitrosomirales* (indicated by the black frame on the genome map rails).

The pan-genome openness of *Nitrososphaerales* was the highest among the four AOA orders (Fig. S3b), and the high genomic diversity was driven by the extensive lateral gene transfer and gene duplication events during *Nitrososphaerales* evolution [12]. The *Nitrosomirales* AOA pan-genome within the available dataset contains a total of 11,334 genes, and the pan-genome graph shows that the sampling of their pan-genome is far from reaching saturation (Fig. S3b). An average of ∼160 novel unique genes per Mbp genome are expected to be identified with each new *Nitrosomirales* species sequenced, which is comparable to the number of new unique genes estimated for *Nitrosopumilales* AOA (Fig. S3c). We further performed Anvi’o pan-genome analysis to assess and visualize *Nitrosomirales*-specific gene contents (Fig. 4). A total of 211,167 coding sequences of the collected AOA genomes (over 50% completeness) were clustered into 13,495 protein clusters. Of these, 1,260 protein clusters were specifically enriched in *Nitrosomirales* genomes but absent or scarce in other AOA orders, including the accessory and unique genes that were assigned to certain Clusters of Orthologous Groups (COGs) functional categories, such as transcription, replication, recombination and repair, energy production and conversion, and transporters as well as many hypothetical genes with unknown functions (Table S7).

Different from the members of their sister order *Ca*. Nitrosocaldales [39, 69, 70], *Nitrosomirales* AOA lacked identifiable hydrogenase genes involved in potential hydrogen oxidation for energy conservation (Table S6). Among the *Nitrosomirales*-specific genes, the genes that encode the major (alpha, containing the active site) subunit of formate dehydrogenase, were found only in *Nitrosomirales* genomes, including those obtained from carbonate and warm springs and deep sandy soils (Fig. 4 and Table S6). *Nitrosomirales* genomes also encodes the putative formate dehydrogenase beta and gamma subunits, which may be involved in electron transfer processes and regulatory functions. In addition, the putative formate transporter genes were identified only in terrestrial *Nitrosomirales* genomes (Fig. 4 and Table S6), strongly suggesting that formate may serve as an alternative electron donor and source of reductant for terrestrial *Nitrosomirales* AOA. It has been shown that *Nitrospira* nitrite-oxidizing bacteria (NOB) species were able to grow using formate as sole substrate with O_2_ or nitrate as terminal electron acceptors [71]. Interestingly, we identified genes that encode the alpha and beta subunits of the putative nitrate reductase in ten terrestrial *Nitrosomirales* genomes (Fig. 4 and Table S6), suggesting that the formate oxidation with nitrate reduction may be directly coupled in *Nitrosomirales* under anoxic conditions, as observed in NOB. Future cultivation and isolation of *Nitrosomirales* AOA are required to confirm their metabolic capacity of using formate as a source of reductant, and such metabolic versatility may enable *Nitrosomirales* AOA to survive during periods of ammonia deprivation.

Other unique genes that are conserved in *Nitrosomirales* AOA include two small blue copper proteins that may function as electron shuttles in the respiratory system, putative CopA and CopD proteins for copper sequestration and uptake, and various regulation proteins that may be associated with stress response and adaptation, such as *Nitrosomirales*-specific transcriptional regulatory and two-component regulatory proteins, DNA repair proteins, and Heat shock proteins (Hsp20 family) (Fig. 4 and Table S6). Putative surface layer (S-layer) protein genes were identified in *Nitrosomirales* genomes (Fig. 4 and Table S6). Moreover, *Nitrosomirales* AOA were found to possess up to three additional putative PKD (polycystic kidney disease) domain protein homologs (Fig. 4 and Table S6), which have been found in extracellular parts of archaeal S-layer proteins that protect cells from extreme environments and assist in cell adhesion and intercellular interaction [72, 73].

### *Nitrosomirales* AOA acquire key metabolic genes via horizontal gene transfer

Similar to other AOA orders, *Nitrosomirales* AOA genomes contain the genes coding for the known ABC subunits of ammonia monooxygenase (AMO), the predicted AMOX, and newly identified AMOY and AMOZ subunits [74] (Fig. 4 and Table S6). The electrons from ammonia oxidation are transferred to oxygen via the copper-centric electron transfer chain, which would lead to the generation of a proton motive force for ATP synthesis via energy-yielding ATPase. The phylogenetic trees of AOA ATPase alpha (A) and epsilon (C) subunits showed clear bifurcating topologies of archaeal-type (A-type) and vacuolar-like (V-type-like) ATPase subgroups (Fig. S4). The A-type ATPase gene clusters were found in both marine and terrestrial *Nitrosomirales* AOA genomes, and their phylogeny tracked organismal phylogeny rather than ecological habitat (Fig. S4). Likewise, the A-type *atp* operon of marine *Nitrosomirales* AOA shared the conserved organization and orientation with those of the terrestrial *Ca*. Nitrosocaldales and *Nitrososphaerales* AOA, but were distinct from marine *Nitrosopumilales* species (Fig. S5). Interestingly, four terrestrial *Nitrosomirales* MAGs encoded the entire gene clusters for both A-type and V-type-like ATPases (Fig. S5). Partial gene clusters of V-type ATPase were also found in the deep-sea MAG MTA5, two additional terrestrial MAGs, and a SAG obtained from groundwaters (Fig. S5). Missing V-ATPase subunits in these *Nitrosomirales* genomes may reflect genome incompleteness.

The V-type ATPase has been proposed to serve to maintain the cytosolic pH homeostasis in acidophilic and acid tolerant AOA as well as deep marine AOA by pumping out excessive cytoplasmic protons under acidic or high-pressure conditions [75]. Recent AOA comparative population genomics and phylogenomic analyses have shown that V-type ATPase genes were widely distributed among deep-sea water column [11] and sedimentary [76] *Nitrosopumilales* AOA and desert *Nitrososphaerales* AOA populations [77]. Apart from the experimentally tested function in low pH adaptation, AOA V-type ATPase may be also coupled to sodium (Na^+^) motive force at high pH levels and protect cells from high-salt stress [24, 77]. The ATPase subunit phylogenetic trees show that the AOA V-type ATPase subgroup rooted with *Nitrosomirales* AOA variants (Fig. S4). Thus, it is most likely that *Nitrosomirales* species also acquired the V-ATPase by horizontal gene transfer, as observed for other AOA order species [75]. It is interesting to note that the V-type *atp* operon was located next to the A-type *atp* operon in *Nitrosomirales* MAGs (Fig. S5). In contrast, although the hadopelagic *Nitrosopumilales* AOA species also contain both types of ATPases, the V-type *atp* operon was distantly located from the A-type *atp* operon (Fig. S5), suggesting the highly mobile V-type *atp* operon may have been reshuffled by several successive events along the evolutionary trajectory of AOA.

Nitric oxide (NO) has been shown as a central intermediate in the archaeal ammonia oxidation pathway [6, 78], and the putative NO-forming nitrite reductase (NirK) is conserved among *Nitrosopumilales* and *Nitrososphaerales* AOA species [11]. However, no gene encoding NirK proteins has yet been identified in *Ca.* Nitrosocaldus AOA species [39, 70]. In contrast, it was identified in thermophilic *Ca.* Nitrosothermus AOA MAGs [69]. NirK homologs were also identified in both marine and terrestrial *Nitrosomirales* genomes, and many *Nitrosomirales* species encode two NirK paralogs in the genomes (Fig. S6). Intriguingly, the *nirK* genes of the deep-sea *Nitrosomirales* MAG MTA5 and warm spring MAG WS3 did not cluster with those encoded by the relatively closely related *Ca*. Nitrosothermus AOA, but rather grouped with those of the distinctly related deep-sea water column B and terrestrial *Nitrosotenuis* AOA, respectively (Fig. S6). This suggests that the *nirK* genes in these *Nitrosomirales* MAGs have been acquired via lateral gene transfer from *Nitrosopumilales* AOA that share similar habitats. In addition, *ureC*, the gene encoding the alpha subunit of urease, was also found in terrestrial *Nitrosomirales* MAGs (Fig. S7) and putatively acquired laterally from other mesophilic terrestrial AOA lineages. The *ureC* genes of *Nitrosomirales* MAGs were phylogenetically distinct from those of thermophilic *Ca.* Nitrosocaldales AOA but nested among *Nitrososphaerales* AOA lineages (Fig. S7). Thus, *Nitrosomirales* AOA may have acquired urea utilization genes to enhance metabolic versatility during their evolution. Taken together, our results highlight the extensive lateral transfer of key genes and operons involved in energy conservation in *Nitrosomirales* AOA, and the acquisition of these essential metabolic genes may have facilitated their radiation into the diversity of subsurface, marine, and geothermal environments they inhabit as now revealed by our metagenomic-based biogeography analysis.

### Hot origin of *Nitrosomirales* and terrestrial-marine habitat transitions within each AOA order

The presence of *Nitrosomirales* sequences in geothermal habitats indicates their broad occurrence in habitats that span a wide range of temperatures (Fig. 2). To estimate the optimal growth temperatures (OGTs) of the ancestors of *Nitrosomirales* AOA and other AOA lineages, we used molecular thermometer-based evolutionary analysis, which involved predicting ancestral 16S rRNA stem composition along the reference tree [43] and using the established linear relationship between the OGTs of cultured AOA species and their 16S rRNA stem GC contents [13, 79]. The predicted ancestral OGT for *Nitrosomirales* AOA was 69 °C (Fig. S8), suggesting that this deeply branching AOA lineage likely originated from hot environments. In addition, the estimated OGTs of the ancestors of *Nitrosopumilales*, *Nitrososphaerales*, and *Ca*. Nitrosocaldales, as well as the last common ancestor of AOA were broadly consistent with the previously reported values by Abby et al. [13] (Fig. S8). Taken together, these results provide additional support for the evolutionary model proposing the hot origin of AOA [13, 80].

Our comprehensive single-gene and whole-genome based phylogenetic analyses with additional *Nitrosomirales* genomes and marine AOA 16S rRNA sequences provide a better resolved framework of the AOA phylogeny and reconstruct a more complex evolutionary trajectory of major AOA lineages than previously proposed. We found that after the initial expansion from hot to moderate temperature environments, AOA diverged via two primary paths of adaptive evolution that ultimately gave rise to the two major contemporary branches: the newly defined *Nitrosomirales* AOA spanning both terrestrial and marine environments, and the previously described mesophilic AOA including *Nitrososphaerales* mostly in terrestrial settings and *Nitrosopumilales* mostly in marine settings (Fig. 1). Previous molecular dating analyses suggested that AOA first transitioned into the temperate terrestrial environments before expanding to marine environments, and further transition from shallow marine sea into deep-sea waters awaited the oxygenation of the deep ocean during the Neoproterozoic [16, 81]. However, we found the presence of thermophilic *Nitrosocaldales* 16S rRNA gene sequences in various high-temperature marine environments, including shallow-sea hydrothermal vents [82], a coastal hot springs (NCBI accession No.: JX047158), deep-sea hydrothermal fields of the Marina Trough [83], and the walls of an active deep-sea sulfide chimney with 302 °C venting liquid [84] (Fig. S9). This strongly suggests the expansion of AOA into both shallow and deep-sea habitats prior to their transition to temperate terrestrial environments and likely places the origin of marine AOA further back in evolutionary history than previously thought. In addition, interestingly, we found *Nitrososphaerales* 16S rRNA sequences were present in many sponge and coral reef samples [62, 63, 64, 85] as well as deep-sea sediments [86] (Fig. S9), revealing the broader habitat range of *Nitrososphaerales* AOA that have been thought mainly restricted to terrestrial environments [22]. Taken together, these results indicate that the terrestrial-marine habitat transitions occurred in not only the order *Nitrosopumilales*, but also each one of the four major orders throughout the evolutionary history of ammonia-oxidizing *Thaumarchaeota* (Fig. S9), depicting a highly dynamic evolutionary process of this globally widespread functional guild.

## Conclusions

Our comparative genomic and phylogenomic analyses of 118 AOA and non-ammonia-oxidizing *Thaumarchaeota* genomes revealed a new AOA cluster, *Ca*. Nitrosomirales, that forms a deeply branching order-level lineage within the class *Nitrososphaeria*. Apart from containing expected gene inventories for ammonia oxidation, carbon dioxide fixation, and B-vitamin biosynthesis, *Nitrosomirales* AOA have the genetic capacity of using formate as an alternative energy source, which may provide metabolic versatility under ammonia starvation. Biogeographic analysis of 16S rRNA and *amoA* genes indicates that *Nitrosomirales* AOA are widely distributed in geothermal, terrestrial, and marine environments, and they are particularly abundant in subsurface terrestrial environments, and some deep-sea sponges. Evidence for independent terrestrial-marine transitions within each AOA order will foster a more detailed understanding of the early evolution and adaptive radiation of archaeal ammonia oxidation during the Proterozoic era.

## Acknowledgement

We thank the DOE Joint Genome Institute for sharing *Nitrosomirales* MAGs and SAGs. This work was supported by the Fundamental Research Funds for the Central Universities of China (to JJ and BW), the National Natural Science Foundation of China grants 41977056 (to BW), 41907027 (to YZ), and the 333 high-level talent project of Jiangsu (to JJ). ML was supported by the National Natural Science Foundation of China grant (32225003, 92251306) and the Innovation Team Project of Universities in Guangdong Province (No. 2020KCXTD023). CZ was supported by the Shenzhen Key Laboratory of Marine Archaea Geo-Omics, Southern University of Science and Technology (ZDSYS201802081843490), and the Southern Marine Science and Engineering Guangdong Laboratory (Guangzhou) (No. K19313901). YZ was supported by the National Natural Science Foundation of China grant 92051114. WMH was supported by Florida Agricultural Experiment Station Hatch project FLA-FTL-005680. WQ was supported by Simons Postdoctoral Fellowship in Marine Microbial Ecology (548565) and the startup funding of the University of Oklahoma.

